# A novel approach to identifying marker genes and estimating the cellular composition of whole blood from gene expression profiles

**DOI:** 10.1101/038794

**Authors:** Casey P. Shannon, Robert Balshaw, Virginia Chen, Zsuzsanna Hollander, Mustafa Toma, Bruce M. McManus, J. Mark FitzGerald, Don D. Sin, Raymond T. Ng, Scott J. Tebbutt

## Abstract

Measuring genome-wide changes in transcript abundance in circulating peripheral whole blood cells is a useful way to study disease pathobiology and may help elucidate biomarkers and molecular mechanisms of disease. The sensitivity and interpretability of analyses carried out in this complex tissue, however, are significantly affected by its dynamic heterogeneity. It is therefore desirable to quantify this heterogeneity, either to account for it or to better model interactions that may be present between the abundance of certain transcripts, some cell types and the indication under study. Accurate enumeration of the many component cell types that make up peripheral whole blood can be costly, however, and may further complicate the sample collection process. Many approaches have been developed to infer the composition of a sample from high-dimensional transcriptomic and, more recently, epigenetic data. These approaches rely on the availability of isolated expression profiles for the cell types to be enumerated. These profiles are platform-specific, suitable datasets are rare, and generating them is expensive. No such dataset exists on the Affymetrix Gene ST platform. We present a freely-available, and open source, multi-response Gaussian model capable of accurately predicting the composition of peripheral whole blood samples from Affymetrix Gene ST expression profiles. This model outperforms other current methods when applied to Gene ST data and could potentially be used to enrich the >10,000 Affymetrix Gene ST blood gene expression profiles currently available on GEO.

**Key Points:**

- We introduce a model that accurately predicts the composition of blood from Affymetrix Gene ST gene expression profiles.
- This model outperforms existing methods when applied to Affymetrix Gene ST expression profiles from blood.

## Introduction

Measuring genome-wide changes in transcript abundance in circulating peripheral whole blood cells is a useful way to study disease pathobiology [1]. By providing a relatively comprehensive survey of the status of the immune system, peripheral whole blood transcript abundances may help elucidate molecular mechanisms [2]. The sensitivity and interpretability of analyses carried out in this tissue, however, are significantly affected by its dynamic heterogeneity [3]. It is therefore desirable to quantify this heterogeneity, either to account for it or to model interactions that may be present between the abundance of certain transcripts, some cell types, and some phenotypic indication.

Accurate enumeration of the many component cell types that make up peripheral whole blood can be costly, however, and may further complicate the sample collection process, beyond a simple complete blood count and leukocyte differentials (CBC/Diffs). Further, the majority of publicly available peripheral whole blood-derived gene expression profiles on the Gene Expression Omnibus [4] do not include any composition information. Accurate quantification of the cellular composition of blood samples from gene expression data without performing additional experiments is useful, allowing for re-analysis of existing public data, for example.

Many approaches have been developed to infer the cellular composition of a sample from high-dimensional transcriptomic [3, 5-10] and, more recently, DNA methylation data [11, 12]. Briefly, if ***X, W,*** and ***H*** are matrices with entries ***X**_ij_* (observed expression for sample ***i***, gene ***j***), ***w**_ik_* (composition for sample ***i***, cell type ***k***), and ***h_kj_*** (cell type-specific contribution to the observed expression for cell type ***k***, gene ***j***), then the problem can be stated: having observed ***X***, we wish to estimate ***W***, based on the assumed relationship between expression and composition:

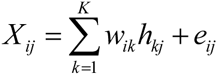

where *e_ij_* represents the expression information for sample *i*, gene *j* that is not predictable by the cell composition.

We further assume that, for each component cell type ***k***, there exists a subset of features ***X^k^_ij_***, in ***X*** whose observed expression in sample ***i*** is proportional to the relative abundance of cell type ***k*** in sample ***i***.

More formally:

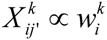

These composition-discriminating features are termed marker genes. For such genes, the elements of the ***H*** can be derived from omics profiles of isolated cell types obtained from reference datasets, and ***W*** estimated by regression [5, 7-13]. Importantly, mapping such marker genes across technology platforms is not always tractable. Not all genes can be readily mapped across gene expression platforms and the values derived from reference datasets may be specific to the platform on which the gene expression was measured. This limits application of these techniques to platforms on which suitable reference datasets exist. Unfortunately, generating such datasets is costly, and they are correspondingly rare.

More recently, so-called reference-free approaches have been proposed to address this issue [6, 14]. When applied to transcriptomic data, these approaches still require the identification of suitable marker genes for the cell types to be quantified. This selection is of paramount importance to achieve optimal performance. The general strategy for marker selection is to identify genes whose expression in one cell type exceeds that of all other cell types being considered [6], a process that itself relies on reference datasets. In fact, all approaches discussed thus far rely on one of a handful of publicly available reference datasets to derive a basis matrix or identify suitable marker genes [12, 15, 16]. No suitable reference dataset exists on the newer Affymetrix Gene ST platform.

Here we propose a new approach that leverages a multi-task learning algorithm to construct a statistical model able to predict the composition of peripheral whole blood from Affymetrix Gene ST expression profiles. We further show that the coefficients of this model can be used to identify suitable marker genes directly, without the need for a reference dataset. Our strategy is readily applicable to other tissues and/or platforms, which would allow for the development of tools to accurately segment and quantify a variety of admixed tissues from their gene expression profiles, to account for cellular heterogeneity across indications or model interactions between gene expression, some cell types and the indication under study. The described model is freely-available and open source, outperforms other current methods when applied to Gene ST data, and could significantly improve our ability to study disease pathobiology in blood by allowing a more complete study of the various components of the immune compartment of blood from whole blood gene expression.

## Patients, material, and methods

### Availability of data and materials

The datasets supporting the conclusions of this article are available on the Gene Expression Omnibus (GEO): repositories GSE77344 (RTP cohort samples) and GSE77343 (CHFP samples). The model is made available as a package for the R statistical programming language, distributed under the GNU General Public License v3.0, and is hosted on GitHub: https://www.github.com/cashoes/enumerateblood.

### Cohorts

We used two large clinical cohorts to train and validate the new statistical model. The Rapid Transition Program (RTP) included prospectively enrolled patients with chronic obstructive pulmonary disease (COPD), presenting either to St. Paul’s Hospital or Vancouver General Hospital (Vancouver, Canada). Subjects presenting to the emergency department or those visiting the COPD clinic were approached for consent to participate in the study. Matched genome-wide transcript abundance and DNA methylation profiles were available for 172 samples from this cohort. This data was used for training the model and cross-validation. Complete blood counts, including leukocyte differentials (CBC/Diffs) were available for all blood samples and used as an independent measure of blood composition (excluding lymphocyte subtypes).

The chronic heart failure (HF) program (CHFP) included prospectively enrolled HF patients presenting to St. Paul’s Hospital or Vancouver General Hospital (Vancouver, Canada). Subjects were approached during their visit to the heart function, pre-transplant, or maintenance clinics, and those who consented were enrolled in the study. A blood sample was collected at the time of enrollment. Genome-wide transcript abundance profiles and complete blood count, including leukocyte differential (CBC/Diffs) were available for 197 HF patients. This data was used to independently validate the performance of the statistical model.

Both studies were approved by the University of British Columbia Clinical Research Ethics Board and Providence Health Care Research Ethics Board and confirm to the principles outlined in the Declaration of Helsinki.

### Sample processing

For all subjects, blood was collected in PAXgene (PreAnalytix, Switzerland) and EDTA tubes. The EDTA blood was spun down (200 x *g* for 10 minutes at room temperature) and the buffy coat aliquoted out. Both PAXgene blood and buffy coat samples were stored at -80°C.

#### Transcript abundance

Total RNA was extracted from PAXgene blood on the QIAcube (Qiagen, Germany), using the PAXgene Blood miRNA kit from PreAnalytix, according to manufacturer’s instructions. Human Gene 1.1 ST array plates (Affymetrix, United States) were used to measure mRNA abundance. This work was carried out at The Scripps Research Institute DNA Array Core Facility (TSRI; La Jolla, CA). The resulting CEL files were processed using the ‘oligo’ R package [17].

#### Dna methylation

For the RTP cohort samples only, DNA was extracted from buffy coat using Qiagen’s QIAamp DNA Blood Mini kits. DNA was bisulphate-converted using the Zymo Research EZ DNA methylation conversion kit, and Infinium HumanMethylation450 BeadChips (Illumina, United States) were used to measure methylation status at >485,000 sites across the genome. This work was carried out at The Centre for

Applied Genomics (TCAG; Toronto, Canada). The resulting IDAT files were processed using the ‘minfi’ R package [18].

### Statistical analysis

Following preprocessing with their respective packages (’oligo’ or ‘minfi’), the normalized data were batch corrected using the ‘ComBat’ algorithm [19], as implemented in the ‘sva’ R package [20].

#### 1. Model training

Next, we inferred the cellular composition of the RTP cohort blood samples from their DNA methylation profiles using the ‘estimateCellCounts’ function provided by ‘minfi’. This function uses publicly available DNA methylation profiles obtained from isolated leukocyte sub-types to infer the relative abundance of granulocytes, monocytes, B, CD4+ T, CD8+ T and NK cells (details in Table 1) with very high accuracy [12, 18]. We compared these composition estimates to those obtained from a hematology analyzer (CBC/Diffs) to assess accuracy.

**Table 1:** Description of predicted leukocytes

| Cell name | Abbreviation Used | Description |
| --- | --- | --- |
| Granulocytes | Gran | CD15+ granulocytes |
| Monocytes | Mono | CD14+ monocytes |
| B cells | Bcell | CD19+ B-lymphocytes |
| T cells (CD4+) | CD4T | CD3+CD4+ T-lymphocytes |
| T cells (CD8+) | CD8T | CD3+CD8+ T-lymphocytes |
| NK | NK | CD56+ Natural Killer (NK) cells |

We then fit a multi-response Gaussian model using elastic net regression via the ‘glmnet’ R package [21] on the genome-wide transcript abundance data, using the DNA methylation-derived cell proportions as response variables. The multi-response Gaussian model family is useful when there are a number of possibly correlated responses – a so called “multi-task learning” problem – as is the case for these cell proportions. Probesets with minimum log_2_ expression < 5.5 across all samples (22,251) were excluded using the ‘exclude’ parameter. We set the elastic net mixing parameter ‘alpha’ at 0.1 to encourage the selection of a smaller subset of genes and chose the regularization parameter ‘lambda’ using the ‘cv.glmnet’ function set to minimize mean squared error (MSE).

#### 2. Estimating out-of-sample performance

Out-of-sample performance of our model was evaluated using 20 x 10-fold cross-validation (not to be confused with the cross-validation performed by ‘cv.glmnet’ in order to choose an effective regularization parameter ‘lambda’). We then validated the accuracy and calibration of our model by comparing its predicted cell proportions to the available CBC/Diffs data in the CHFP cohort. Unfortunately, a more complete enumeration of the lymphocyte compartment (e.g., by flow cytometry) was not available in any of our cohorts, so we could not independently validate performance in the various lymphocyte sub-types. Instead, the sum of the predicted B, CD4+ T, CD8+ T and NK cell proportions was compared to total lymphocyte proportions from the CBC/Diffs.

#### 3. Performance compared to other current approaches

Finally, we compared the performance of our model to two alternative approaches for determining the composition of blood samples from their gene expression profiles, described by Abbas *et al.[5]* and Chikina *et al.* [6], in this independent heart failure cohort. First, the basis matrix from Abbas *et al.,* derived from the IRIS (Immune Response In Silico) reference dataset, was used to predict the cell proportions of neutrophils, monocytes, B, CD4+ T, CD8+ T and NK cells [15]. Again, the Abbas predicted proportions for B, CD4+ T, CD8+ T and NK cells were summed to obtain a predicted lymphocyte proportion. The Abbas predicted neutrophil, monocyte and lymphocyte proportions were compared to CBC/Diffs.

#### 4. Model features as marker genes for use with reference-free approaches

Next, we evaluated whether our approach could be used to identify more suitable marker gene sets compared to a reference dataset approach. The reference-free approach described by Chikina *et al.* does not require a basis matrix, relying instead on a set of putative marker genes. These are used to guide the decomposition of the dataset’s covariance structure into separate variance components, using singular value decomposition (SVD). Marker genes for each cell type are summarized in this manner, a technique known as eigengene summarization [22]. Given a good set of marker genes, these summarized values, termed surrogate proportion variables, should track with mixture proportions. We used the reference-free approach described by Chikina *et al.* (as implemented in the ‘CellCODE’ R package) and marker genes derived either from the IRIS reference dataset, as recommended by Chikina *et al.,* or from the coefficients of our model. We then compared the surrogate proportion variables produced by ‘CellCODE’, using either marker gene sets, to those obtained from CBC/Diffs in order to see whether we could identify better marker genes. Spearman’s rank correlation (ρ) was used to summarize association between predictions and root mean squared error of prediction (rmse) was used to summarize accuracy and precision.

## Results

DNA methylation derived predictions of the cellular composition of the RTP cohort blood samples were accurate when compared to those obtained from CBC/Diffs (root mean squared error [rmse] = 0.01 -0.08, Spearman’s ρ = 0.85 - 0.94; Supplementary Figure S1). The observed error rates were consistent with those previously reported [11, 12]. These predictions were used as the response variables in a multi-response Gaussian model fit to the RTP cohort gene expression data using an elastic net regression. The model selected by ‘cv.glmnet’ retained 491 features. Its fit to the data is visualized in Figure 2, against both the DNA methylation derived composition estimates (Figure 2A), and CBC/Diffs (Figure 2B). Model fit was good (DNA methylation composition: rmse = 0.01 to 0.04; p = 0.86 to 0.97; CBC/Diffs: rmse = 0.01 to 0.06; p = 0.91 to 0.97) across all cell types, with the exception, perhaps, of CD8+ T cells. When considering the model fit to the CBC/Diffs data, we noted slight bias, with granulocyte proportions tending to be under-predicted and lymphocyte proportions over-predicted.

**Figure 1:**
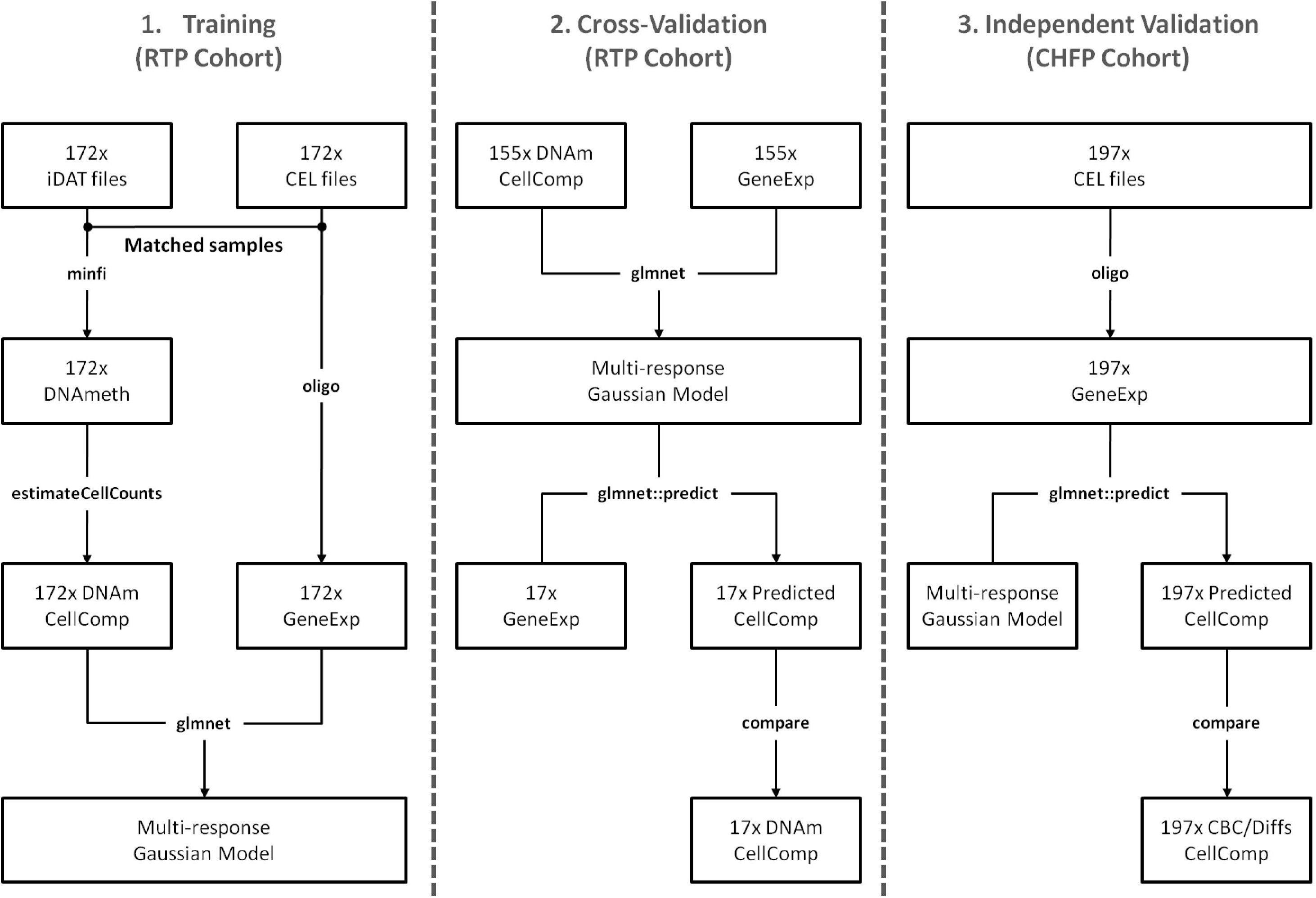
Schematic representation of the experiment. The model was trained using 172 blood gene expression profiles from the Rapid Transition Program cohort (RTP). For all training samples, cellular composition was first estimated from their DNA methylation profiles (using minfi’s ‘estimateCellCounts’) and then used as the response matrix to fit a multi-response Gaussian model (using glmnet) on the blood gene expression profiles (1). The performance of this model on new data was estimated using cross-validation (2) and confirmed using 192 blood expression profiles from the Chronic Heart Failure Program (CHFP), an independent test cohort (3).

**Figure 2:**
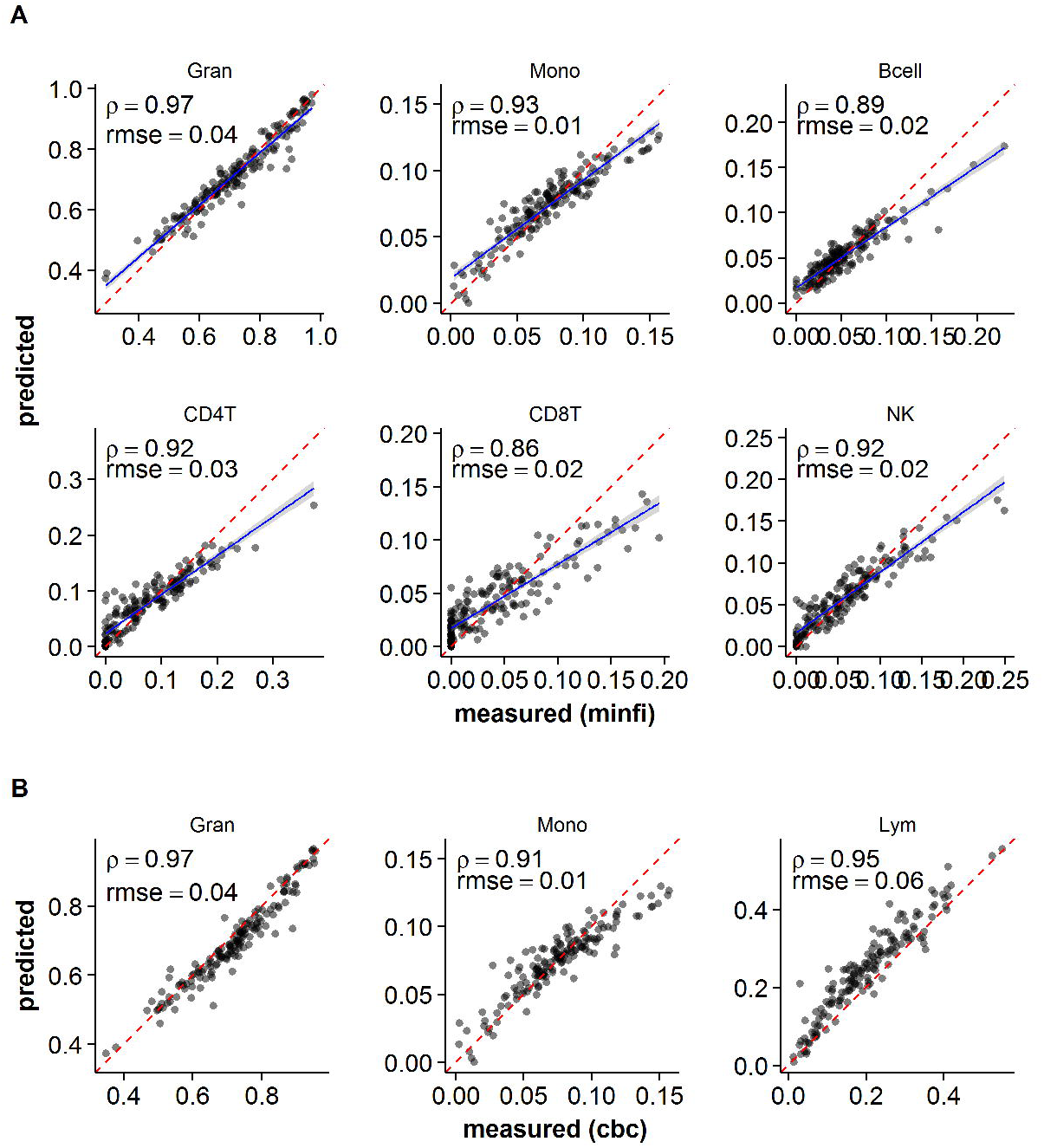
Assessing model fit. Predicted proportions from the model are plotted against the DNA methylation-derived cell proportions for each sample in the training data (**A**) or that obtained from CBC/Diffs (**B**). For **A**, linear best-fit line to the data is plotted (blue line) with 95% point-wise confidence interval for fit (grey band) and compared with perfect agreement (red dashed line). For **B**, the sum of the predicted B, CD4+ T, CD8+ T and NK cell proportions is compared to the total lymphocyte proportions from the CBC/Diffs. For each cell type, Spearman’s rank correlation (ρ) and the root mean squared error (rmse) are reported.

To characterize the potential performance of this model on new data, we carried out a 20 x 10-fold cross-validation. Estimated out-of-sample performance varied across cell types (Figure 3). We report the mean rmse (scaled to the expected cell abundance) and Spearman’s ρ across all 200 generated models. Scaled rmse was lowest for granulocytes (0.08) and monocytes (0.24), higher in B, NK and CD4+ T cells (0.51, 0.52, and 0.58, respectively), and highest in CD8+ T cells (1.21). Absolute rmse (0.02 - 0.06) compared favorably to other methods for inferring cellular composition of samples from gene expression data [5, 7, 8, 13]. Results for Spearman’s ρ were consistent: highest in granulocytes (0.926), followed by monocytes (0.824), NK cells (0.812), CD4+ T cells (0.785), B cells (0.731), and CD8+ T cells (0.671).

**Figure 3:**
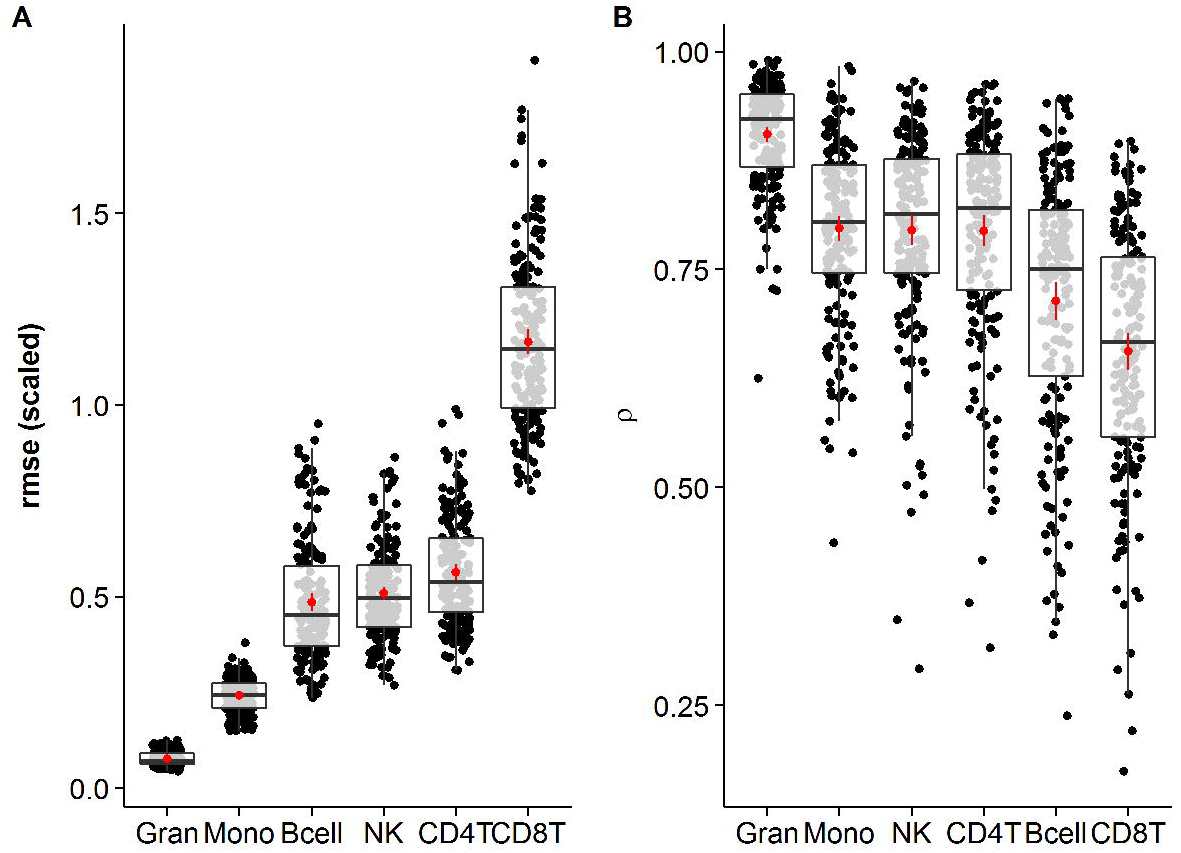
Cross-validation performance. Distribution of root mean square error (rmse; **A**), scaled to the expected frequency for each cell, and Spearman’s rank correlation (p; **B**) for out-of-sample predictions across a 20 x 10 fold cross-validation are visualized using box-and-whisker plots. The mean and 95% CI are shown as a point and range in the center of each boxplot and represent the expected out-of-sample performance.

Next, we applied the model to gene expression profiles from the CHFP cohort blood samples in order to independently validate the model’s performance. Performance remained good (rmse = 0.02 to 0.09; ρ = 0.69 to 0.91; Figure 4A), though the bias we previously noted was more pronounced. Prediction of monocyte proportions was significantly worse than that seen in-sample (ρ = 0.69 vs. 0.91) and expected out-of-sample (from cross-validation; ρ = 0.80 vs. 0.91). Comparing performance of this model against another available approach for inferring the composition of whole blood samples from microarray gene expression data [5], we find that our model performs better, with both correlation to CBC/Diffs data and prediction error markedly improved, especially for monocytes (lymphocyte rmseAbbas= 0.28 vs. rmseglmnet = 0.09; monocyte rmse_Abbas_= 0.07 vs. rmse_glmnet_ = 0.02; ρ_Abbas_ = 0.31 vs. ρ_glmnet_ = 0.69; Figure 4B).

**Figure 4:**
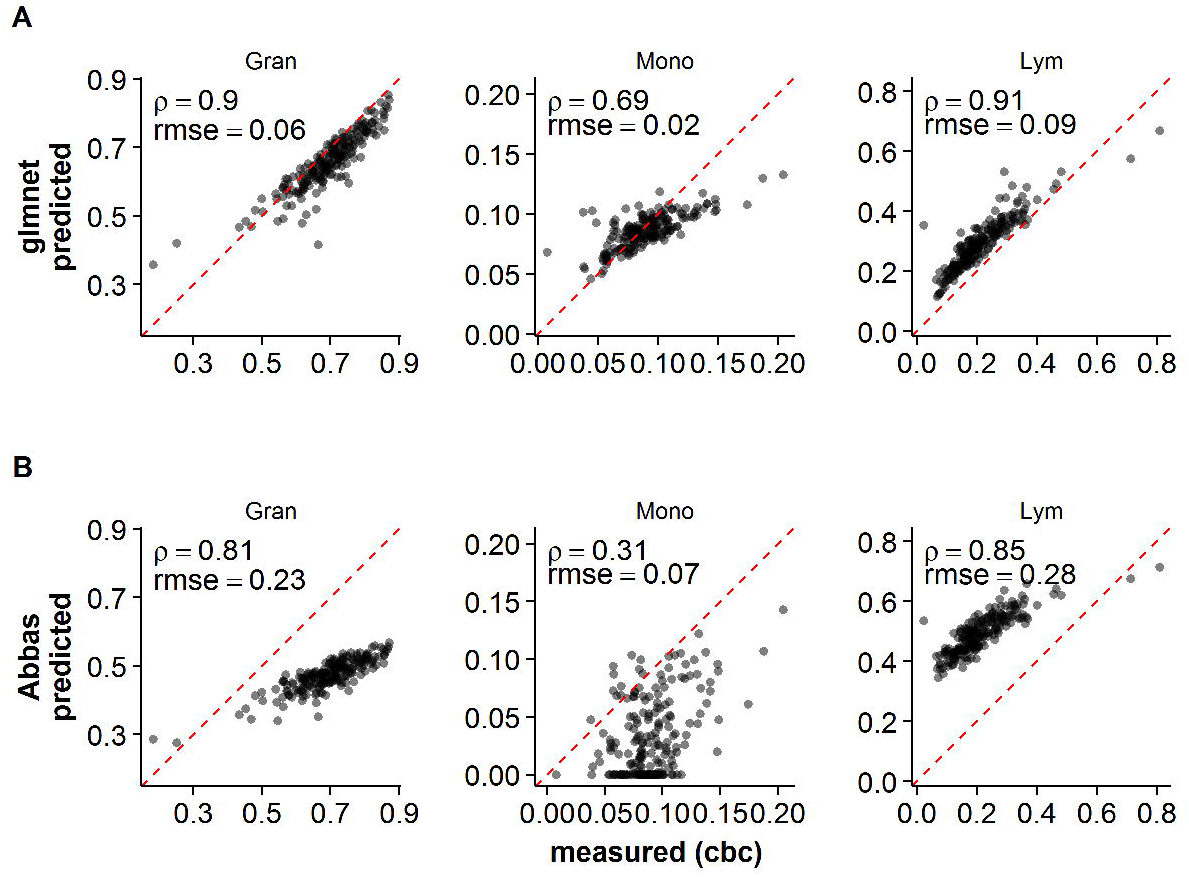
Our model accurately predicts the cellular composition of blood samples and outperforms existing approaches in Affymetrix Gene ST data. Predicted cell proportions are plotted against the cell proportions obtained from CBC/Diffs in an independent dataset (CHFP cohort) for the model **(A)** or using the method from Abbas *et al.* **(B)**. The sum of the predicted B, CD4+ T, CD8+ T and NK cell proportions is compared to the total lymphocyte proportions from the CBC/Diffs. For each cell type, Spearman’s rank correlation (p) and the root mean squared error (rmse) are reported.

Marker genes derived from the coefficients of our model outperformed those derived from the IRIS reference dataset when used to predict cellular composition using the approach proposed by Chikina *et al.* (granulocytes ρ = 0.87 vs. 0.67, lymphocytes ρ = 0.84 vs. 0.78, and monocytes ρ = 0.73 vs. 0.32; Figure 5). The marker gene sets showed minimal overlap (granulocytes = 3/51, monocytes = 4/58, B cells = 0/55, CD4+ T cells = 0/11, CD8+ T cells = 1/15, NK cells = 6/22).

Finally, we applied the model to predict the composition of the RTP cohort blood samples from their gene expression. This is a contrived example, as this information was already available to us, but it serves to illustrate a possible application of the approach. As expected, large differences exist in the proportions of the various cell types between patients given prednisone or not. Patients on prednisone had proportionally lower amounts of monocytes (p = 2.9 x 10^−4^), B (p = 6.6 × 10^−5^), CD4+ T (p = 6.6 × 10^−7^), CD8+ (p = 5.0 × 10^−10^) and NK cells (p = 9.3 × 10^−10^), and proportionally higher amounts of granulocytes (p = 2.3 × 10^−8^).

**Figure 5:**
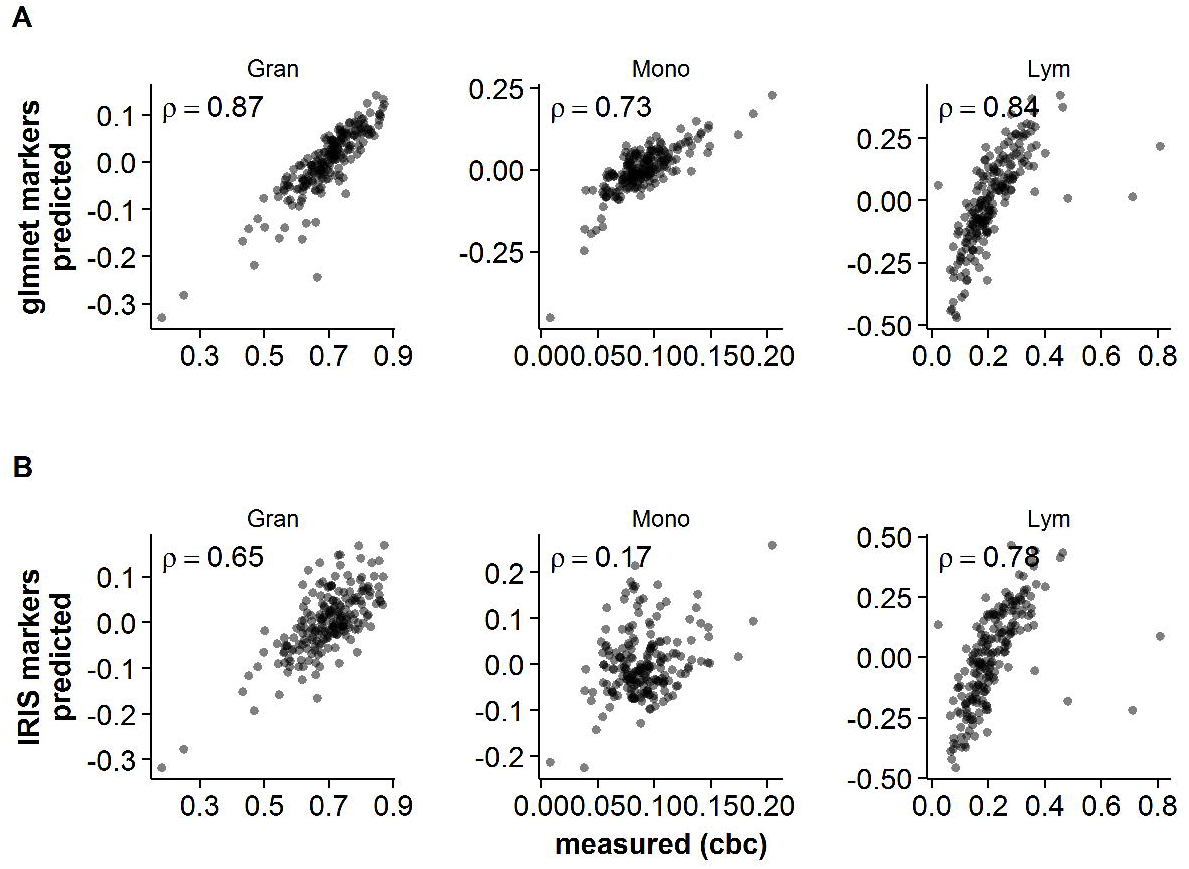
Our model identifies better performing marker genes for use with reference-free approaches in Affymetrix Gene ST data. Surrogate proportion variables obtained from CellCODE are plotted against the cell proportions obtained from CBC/Diffs in an independent dataset (CHFP cohort). The sum of the surrogate proportion variables obtained for B, CD4+ T, CD8+ T and NK cells is compared to the total lymphocyte proportions from the CBC/Diffs. Marker genes used by CellCODE were derived from the coefficients of the model (**A**) or using the recommended set of marker genes (**B**) derived from the IRIS reference dataset. For each cell type, Spearman’s rank correlation (ρ) is reported.

**Figure 6:**
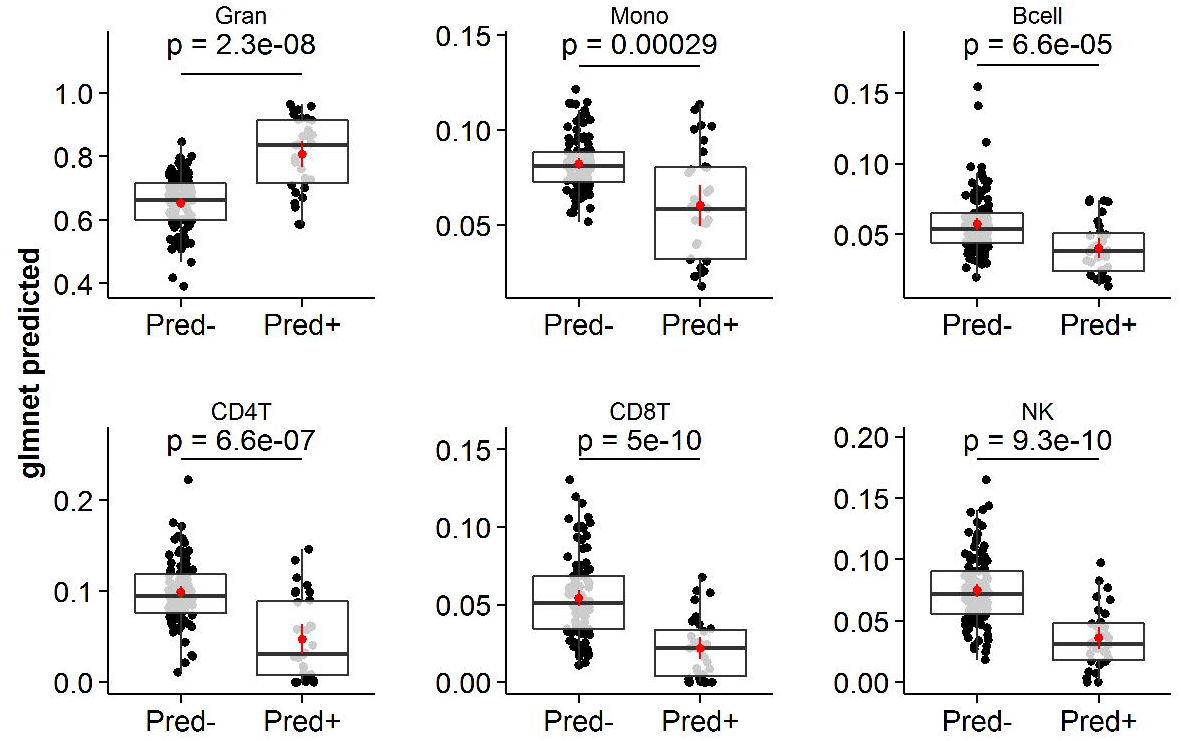
Model predicted cell proportions highlight prednisone-dependent changes in peripheral blood composition. Treatment of acute exacerbations (AE) in COPD with prednisone results in important changes in the cellular composition of peripheral blood. The distributions of granulocyte, monocyte, B, CD4+ T, CD8+ T and NK cell proportions are visualized for patients from the Rapid Transition Program (RTP) cohort that were given prednisone or not (p-value is for the unpaired Student’s t-test comparing the two groups in each case).

## Discussion

We introduce a statistical model for predicting the composition of blood samples from Affymetrix Gene ST gene expression profiles. We demonstrate that this model has suitable performance across all included cell types in cross-validation, and validate its performance in an independent cohort. The training and validation cohorts represent 2 major clinical indications, COPD and CHF, and include patients with various comorbidities, on various drugs, some with strong effects on blood gene expression (e.g., prednisone), suggesting that our model may generalize well and be broadly applicable. All training and validation samples were from older individuals, however, and it may be that this model will not generalize well to pediatric populations. A loss of performance in pediatric population has been noted when using a similar approach with DNA methylation data [23].

We also show that platform-specific marker gene sets can be derived without the need for reference datasets of isolated gene expression profiles for the cell types we wish to enumerate. Using marker genes selected from the coefficients of our model in combination with the reference-free approach proposed by Chikina *et al.* resulted in better performance compared to using marker genes derived from isolated leukocyte gene expression profiles obtained on another microarray platform. Interestingly, the reference-free approach performed only slightly worse than our model, although with loss of scale. This suggests that the non-zero coefficient weights of the model (which we used to select marker genes for the various included cell types) can be estimated entirely in the data, and that these marker genes may be context-independent surrogates of cell proportions.

More generally, the strategy we adopted to derive our model (and identify suitable marker genes) could be readily applied to other platforms, or tissues of interest. The only requirements are accurate quantification of the cell types of interest across a large cohort with matched omics profiling. For many popular platforms (e.g., RNA-seq), this schema may be more cost effective than sorting and profiling a number of replicates for all cells of interest, particularly when we consider how costs would scale with additional cell types to be quantified. Moreover, for low abundance cell types, obtaining a sufficient quantity to profile may not be feasible, depending on the efficiency of available separation techniques, amount of admixed tissue that can be collected in practice.

The lack of independent validation within the lymphocyte sub-types is a limitation, though cross-validation performance was good across all cell types. We believe it is unlikely that poor performance in some or all lymphocyte sub-types would result in good performance when summed and compared to CBC/Diffs. Model fit exhibits some degree of shrinkage (flattening of the plot of predicted vs. observed away from the 45 degree line). This is expected, however, and related to the phenomenon of regression to the mean. Performance in cross-validation was notably worse for CD8+ T cells. This could be because of the preponderance of zero values for this particular cell type. We also note that performance in monocytes drops significantly in the validation cohort. It is unclear why this is, but one possibility is the difference in the distribution of values in the validation cohort (mean monocyte proportion in training: 0.073 vs. 0.090 in the validation; p = 1.39 x 10^−7^). We have observed poor performance of various deconvolution approaches in quantifying monocytes in the past [13, 24]. It might be that circulating monocyte diversity is poorly reflected in our current framework and we may be selecting poor marker genes for this cell type as a result. A similar rationale could be applied to explain the poor CD8+ T cell performance results in cross-validation. Certainly, it offers the opportunity for further exploration of the true complexity of these cell types in peripheral blood.

In summary, our freely-available, open source statistical model is capable of accurately inferring the composition of peripheral whole blood samples from Affymetrix Gene ST expression profiles. The strategy we adopted to derive this model is readily applicable to other tissues and/or platforms, which would allow for the development of tools to accurately segment and quantify a variety of admixed tissues from their gene expression profiles, to account for cellular heterogeneity across indications or model interactions between gene expression, some cell types and the indication under study. The described model outperforms other current methods when applied to Gene ST data and significantly improves our ability to study disease pathobiology in blood. We provide the opportunity to enrich the >10,000 Affymetrix Gene ST blood gene expression profiles currently available on GEO, by allowing a more complete study of the various components of the immune compartment of blood from whole blood gene expression.

## Acknowledgements

The authors would like to thank the research participants without whose tissue donations none of this work would be possible. Additional thanks to study nurses and clinical coordinators for their contributions to patient recruitment and data collection. The authors would also like to acknowledge Dr. Karen Lam for her insightful comments and discussion.

Funding agencies: Genome Canada, Genome British Columbia, Genome Quebec, Canadian Institutes of Health Research, Providence Health Care, St. Paul’s Hospital Foundation, and PROOF Centre

## Authorship and Conflict-of-Interest Statements

Collected data: MT, BMM, JMF, and DDS. Designed the research: CPS, RB, RTN, and SJT. Analyzed and interpreted data: CPS, VC, and ZH. Performed the statistical analysis: CPS. Wrote the manuscript: CPS. The authors declare no conflict of interest.

---

**Supplementary Figure S1:**
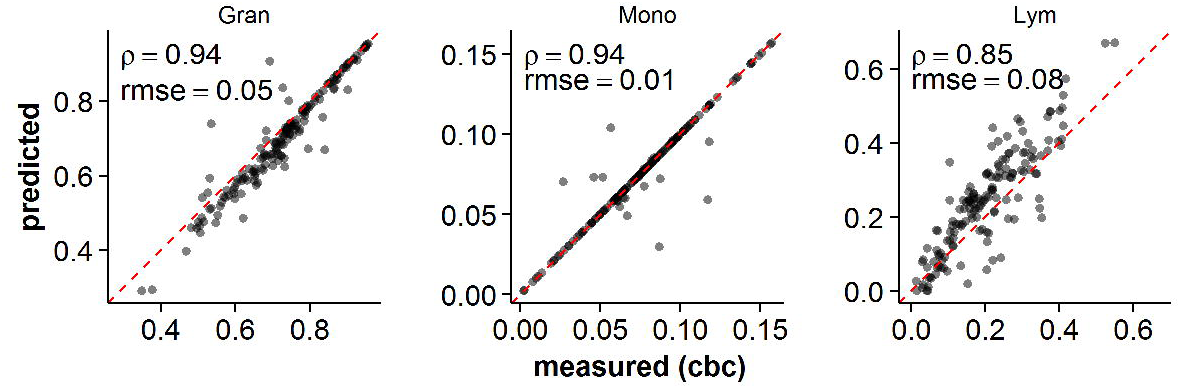
DNA methylation-derived composition vs. CBC/Diffs. Predicted proportions were obtained by applying the ‘estimateCellCounts’ function from the ‘minif R package to peripheral blood derived DNA methylation profiles in the Rapid Transition Program (RTP) cohort and plotted against cell proportions obtained from CBC/Diffs. The sum of the predicted B, CD4+ T, CD8+ T and NK cell proportions is compared to the total lymphocyte proportions from the CBC/Diffs. For each cell type, Spearman’s rank correlation (ρ) and the root mean squared error (rmse) are reported.

